# Small circuit principles for learning in large worlds

**DOI:** 10.1101/2025.11.13.688216

**Authors:** Dhruva V. Raman, Christopher R. Dunne, Katie Davyson, Timothy O’Leary

## Abstract

Animals must learn and make predictions in a world that is not only uncertain, but shaped by unmodelled contingencies: ‘unknown unknowns’. Despite this, current theories of how this is achieved are based on optimal (Bayesian) strategies in simple settings, where it is possible to fully represent a model of the world and infer its state based on sensory data. This could be impractical in situations where an animal has limited prior experience and cognitive capacity, particularly in short-lived animals with small brains. Using the neural circuitry of Drosophila as inspiration, we derive a learning framework, which we call ‘Hetlearn’, for coping in such situations. Hetlearn sacrifices asymptotic optimality on idealised inference tasks, making it less fragile to abrupt changes in environmental statistics and providing more accurate state information than standard Bayesian procedures when data are limited. Hetlearn explains key physiological and anatomical features of the Drosophila Mushroom body (MB), and predicts behavioural features of learning that are conserved across species, such as second-order conditioning and reversal learning.

## Introduction

Animals learn from experience to predict future events. Existing theory frames this as state inference: updating a probabilistic map of relevant aspects of the environment, called *states*, and predicting how these states evolve as a function of time, context, actions and external events [35, 7, 15]. Simple, familiar situations such as repeated laboratory tasks can provide a mathematically tractable environment where an agent can represent all possible states, actions, and consequences, allowing for optimal probabilistic reasoning [14, 19, 31].

Despite their utility in establishing normative principles for reasoning and decision making, idealised tasks whose causal structure can be fully modelled and learned occupy what is known as a ‘Small World’ [43]. This contrasts with the real world, or ‘Large World’: a messy reality of unforeseen contingencies (“unknown unknowns”), and open-ended state spaces where optimal inference is intractable.

Animal brains operate in the Large World setting [29]: they inhabit the real world, and even simpler, restricted situations (such as laboratory tasks) may present animals with too little data to perform optimal inference. Under these conditions, an optimal Small World learning procedure may be less relevant for understanding biological learning, particularly if the optimal procedure is fragile to violations in modelling assumptions [22].

Using well-established features of Drosophila learning circuitry and behaviour, we devised a novel learning algorithm ‘Hetlearn’ (for heterologous learners) designed for rapid, approximate inferences that are robust to unmodelled causes. Hetlearn incorporates a key feature of the mushroom body: parallel, associative learning modules that operate independently on fast and slow timescales. This feature addresses a well-known issue with Bayesian inference by effectively mitigating ambiguities between stochasticity and volatility [6, 41]. Remarkably, in a low to moderate data regime of hundreds of samples, Hetlearn outperforms asymptotically optimal, Bayesian algorithms despite being suboptimal in the long run. This regime is likely relevant to animals making decisions with little data in volatile environments.

Hetlearn recreates Drosophila behaviour across several chosen tasks: second-order, extinction, and reversal conditioning. It explains both behaviour and the heterologous dynamics of dopaminergic signalling in the mushroom body. Furthermore, it provides new, testable hypotheses on the roles of poorly-understood inter-compartmental mushroom body interactions during these tasks. More abstractly, it provides a normative principle for learning in *truly* novel environments where there is uncertainty in the values of represented states (as in probabilistic inference), but also uncertainty in which states to represent in a cognitive map.

## Results

### State inference in a Large World

We first motivate Hetlearn abstractly, using the standard probabilistic inference framework as a comparison. In doing so, we will expose two well-known issues with probabilistic inference which can be detrimental to biological learning in naturalistic settings, and that Hetlearn addresses.

A learner undertaking probabilistic state inference at time *t* holds and updates a probability distribution *P* [*x*_*t*_, *θ* | *y*_1:*t*_] on dynamic states *x*_*t*_ and static parameters *θ* that collectively represent the environment, given an observation history *y*_1:*t*_. The optimal update for this distribution uses Bayes’ rule, which requires a pre-specified generative model (or cognitive map [5]) of the environment comprising probability distributions for state evolution: *P*_*x*_(*x*_*t*_|*x*_*t*−1_, *θ*), and observation: *P*_*y*_(*y*_*t*_ |*x*_*t*_, *θ*). This generative model is variously assumed to be learned from a mixture of prior experience and developmental processes that embed innate assumptions about the world.

Generative models provide probabilistic outcomes. This is assumed to account for the effects of countless unmodelled ‘large-world’ factors *z*_*t*_, which collectively make the future unpredictable from the perspective of the learner. Specifically, we could specify ground-truth models of state update and observation as *x*_*t*_ = *f* (*x*_*t*−1_, *z*_*t*_), and *y*_*t*_ = *g*(*x*_*t*_, *z*_*t*_). Probabilistic inference assumes that

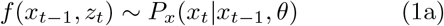

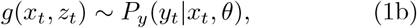

where the distributions *P*_*x*_ and *P*_*y*_ are known ahead of time as part of the generative model, and the parameters *θ* are learnt during the task.

This theoretical framework raises two profound questions that form the theoretical motivation for Hetlearn. First, even if the generative model specified in Eqs. 1 accurately reflects ground-truth, its use in state estimation may be inherently fragile to approximation errors, particularly in situations where data to calibrate states is limited. This is inevitable for algorithms implemented in biological circuits, and could leave an animal with a predictive model that is both algorithmically complicated and poorly performing. There may be algorithmic tricks by which an inherently suboptimal, simpler inference can yield predictions that are more accurate in the short-term. Hetlearn implements these tricks.

Second, systematic ‘large-world’ changes in the environment can fundamentally invalidate the animal’s internal generative model. If the true data-generating process shifts, the animal’s cognitive map and prior assumptions are decoupled from reality. Consequently, the theoretical guarantees underpinning the reliability of Bayesian inference (such as the evidence lower bound [52]) cease to apply, as they remain tied to an outdated representation of the world. Again, there is no question that animals can fail to adapt their behaviour to new ecological niches. However, a simple algorithmic means of minimising the fragility of an animal’s predictive model to Large World effects would clearly offer a reproductive advantage. Hetlearn offers a means of achieving this.

We will next construct and analyse a simple example that illustrates how Hetlearn addresses the first issue by outperforming a Bayesian inference procedure in the low-data regime. Later sections will tackle abrupt changes to generative models.

### Basic Hetlearn algorithm for learning via prediction errors

Many models of learning posit that animals update state estimates from prediction errors (PEs) [45]. States are taken to represent physical or abstract variables, from properties of visual stimuli to reward. Notable examples of learning algorithms that are proposed to model biological state estimation are Kalman filtering [11, 25, 20] and temporal difference reinforcement learning [44, 46, 39, 9]. A common dilemma faced by any learning algorithm is that there is a tradeoff in deciding to what extent a PE is attributable to stochastic fluctuations that should be ignored, versus systematic changes (often known as volatility) that should be adapted to.

We first propose a simple algorithm for updating to PEs and claim it has advantages over classical probabilistic inference in biologically relevant scenarios: low data, limited algorithmic resources. Our setting is a learner attempting to infer a single state from ambiguous observations over repeated trials. For concreteness, we consider the state as a reward 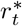 associated with that state. The received reward at time *t, r*_*t*_, takes the form:

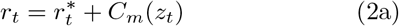

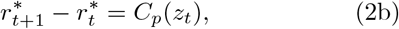

This models a general situation where large-world causes of systematic change (volatility, *C*_*p*_(*z*_*t*_)) and unsystematic, noisy fluctuations (stochasticity, *C*_*m*_(*z*_*t*_)) both affect *r*_*t*_. The learner holds a state estimate 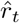, with estimation error: 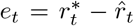. This estimate is updated from prediction errors: 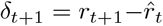.

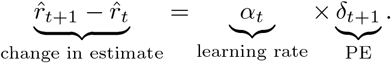

An excessive learning rate (*α*_*t*_ ≈1) erases valuable accumulated experience by overreacting to stochastic fluctuations. An insufficient learning rate (*α*_*t*_ ≈ 0) cannot keep up with systematic volatility, or reduce a large estimation error. The truly optimal learning rate (zero estimation error on trial *t* + 1, see SI S1) is

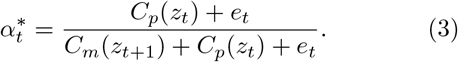

This learning rate in inaccessible to any learning algorithm: *C*_*m*_, *C*_*p*_, and *e*_*t*_ will fluctuate on each timestep, and their values cannot be disambiguated from the scalar PE signal which sums their individual effects.

To deal with this, and in agreement with neural circuit properties that we discuss in detail later, we propose the learner maintains *multiple* parallel valence estimates (sublearners), each with intrinsic, fixed guesses of *α*^*^ within extremal values of 0 and 1. To form an estimate, the learner weights the sublearners according to their recent predictive performance.

### A Hetlearn strategy for learning from prediction errors

We formulate a simple Hetlearn strategy for PE updates: Take *n* parallel sublearners, each with their own estimates: 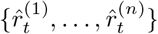. The *i*^*th*^ learner receives a prediction error 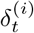, and updates its estimate as:

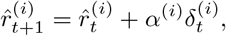

where *α*^(*i*)^ is fixed and heterogeneous across sublearners. The overall prediction is a weighted vote: 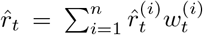, where 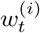 is the weight assigned to the *i*^*th*^ sublearner.

To determine the weights, the agent uses an estimate of recent error for each sublearner. One simple estimate is:

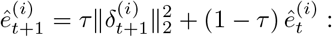

a moving average of PE magnitude, with recency bias 0 ≤ *τ* ≤ 1. Weights are then 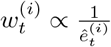, normalised so that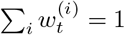. The free parameters in this strategy are the learning rates *α*^(*i*)^ of each sublearner, and the recency bias *τ* .

Hetlearn differs importantly from probabilistic inference, where large-world effects must be modelled as probability distributions to infer learning rate (Eq. (1)). We now demonstrate this on a simple didactic example, where we compare Hetlearn to the Kalman Filter (KF): a popular model for how animals learn from prediction errors [11, 25, 20], with biologically plausible neural circuit implementations [12, 54]. The KF assumes fixed Gaussian distributions:

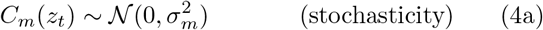

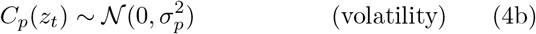

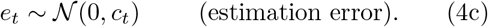

With these assumptions, baseline reward is a Gaussian distribution whose mean updates with a dynamic, Bayes-optimal learning rate. The learning rate calculation, however, requires knowledge of 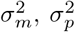, and *c*_0_ as part of its generative model (Methods, (7)). We can say that these form part of the fixed ‘small-world’ structure assumed by the model.

In Figure 1, we simulate a large-world where 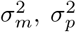, and *c*_0_ change abruptly, halfway through the simulation (trial 500). The KF is seeded with initially correct values for these parameters. Hetlearn doesn’t require or use knowledge of these parameters, and is thus slightly suboptimal in the first half of the simulation. This parsimony makes it more robust to the abrupt large-world change, to the extent that its cumulative error over the entire simulation is markedly lower than the KF.

**Figure 1.**
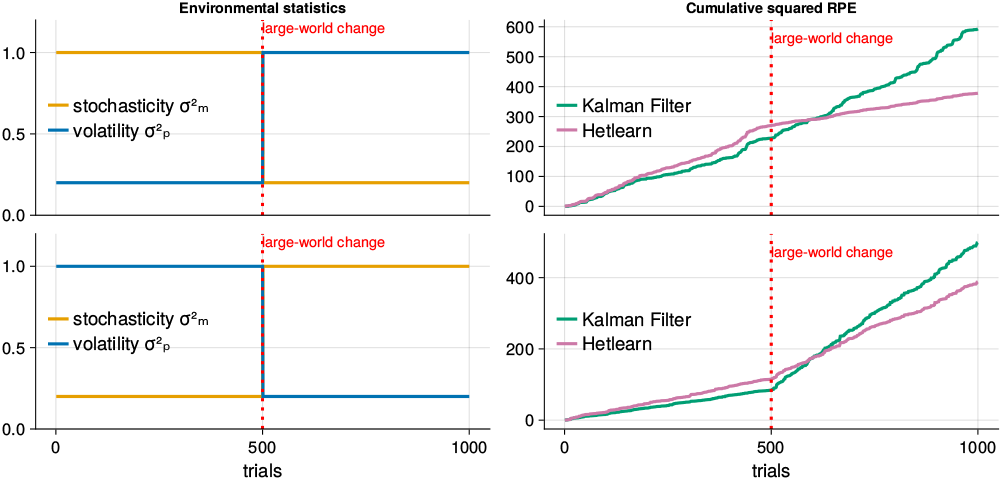
The Kalman Filter requires seeding with values for reward stochasticity/volatility. It is seeded with the initial values, for which it is statistically optimal. When the values change due to unmodelled, unrepresentable large-world effects, it performs poorly. A Bayes-optimal learner can be *fragile* to large-world effects that break its foundational assumptions. Hetlearn is suboptimal for any particular, fixed reward stochasticity/volatility, but robust to changes.

More generally, any KF requires direct access to the correct stochasticity/volatility values, which stretches the plausibility of a biological implementation. Consequently, studies that propose the KF as a model of biological learning often hardcode guesses into a model [11, 12, 20]. Figure 1 suggests that a nonoptimal, non-probabilistic strategy could be preferable where such guesses, which form part of the generative model structure, are unreliable or subject to change.

The standard KF deployed in Figure 1 could be improved by incorporating stochasticity and volatility as parameters to be concurrently estimated. More generally, generative models can be improved by adding flexibility through more parameters. Previous work has thus proposed algorithms that learn stochasticity and volatility online [6, 41]. This clearly results in greater algorithmic complexity. More importantly, we claim that it necessarily results in worse short-term performance than Hetlearn.

To demonstrate, we first implemented an (augmented) Kalman learner that additionally estimates 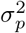 and 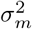 from historical rewards, using the state-of-the-art Autocovariance Least Squares (ALS) method [38]. ALS regresses point-estimates of 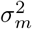 and 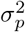from the history of reward prediction errors (see Methods). The usage of ALS means the learner is undertaking approximate Bayesian inference: updating complete probability distributions of stochasticity and volatility Bayes-optimally is analytically intractable. Meanwhile we provide *c*_0_ explicitly to the learner.

The performance of the Kalman learner is shown in Figure 2. For over 100 trials, Hetlearn markedly outperforms in mean-square prediction error, with cumulative error lower for Hetlearn until past the 500th trial. Hetlearn superiority is also robust to choices of the (fixed) recency bias parameter, *τ* (see SI S5). In the long run, as expected, the Kalman learner converges on the true stochasticity/volatility values (Figure 2A,C), and asymptotically approaches Bayes-optimal performance.

**Figure 2.**
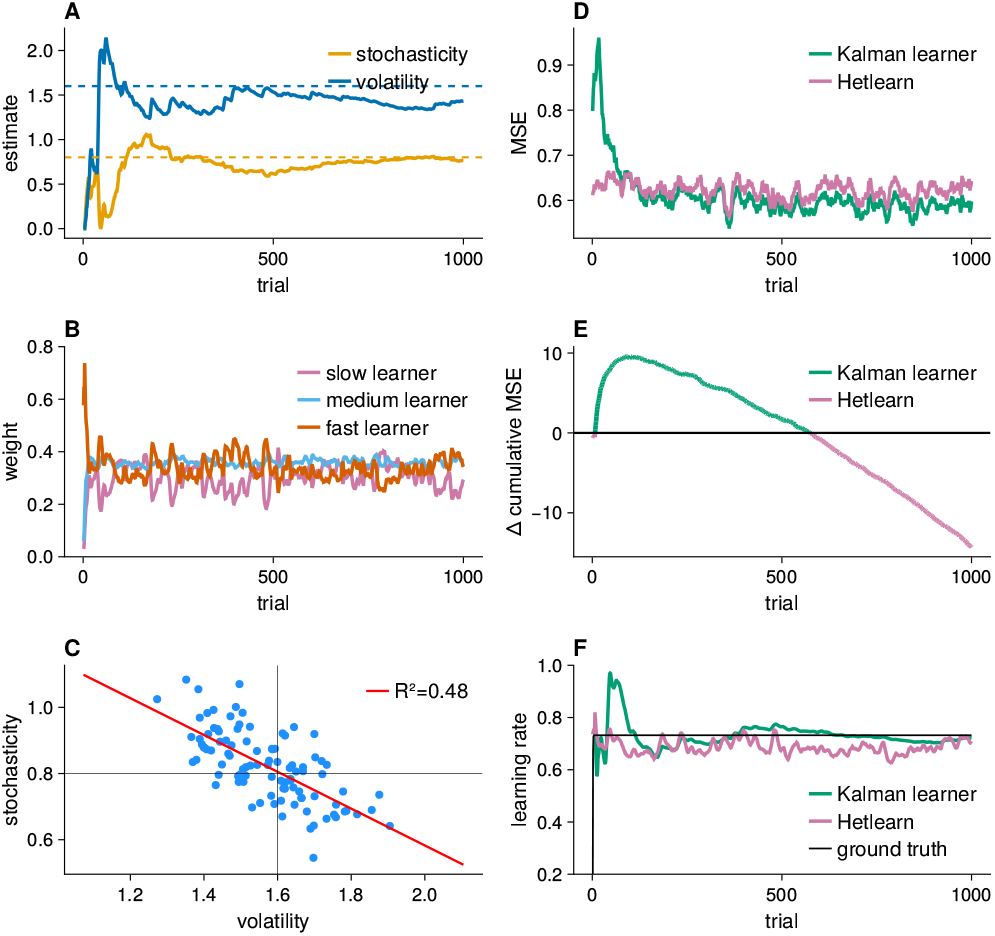
**A**: Convergence of ALS algorithm’s stochasticity and volatility estimates over time, compared to their (dotted) true values. One run plotted for illustration. **B**: Faster convergence of the weightings of Hetlearn sublearners over time. One run plotted for illustration **C**: 100 samples of the final ALS estimates of stochasticity and volatility, after 1000 timesteps. Estimation errors are anticorrelated. **D**: Mean MSE of Kalman learner vs Hetlearn over 100 repeats. Bands represent standard error. **E**: Mean trajectory (100 repeats) of the difference in cumulative MSE for the Kalman learner and Hetlearn. Green (positive) values indicate that the Kalman learner had higher cumulative MSE than Hetlearn, while purple (negative) values indicate the opposite. **F**: Learning rate of the two algorithms as compared to the optimal formula of Eq. (3) over a single trial.

Hetlearn possesses a key property that provides robustness to changing conditions and limited data. The (expected) estimated error of a sublearner with fixed learning rate *α* converges to its asymptotic value 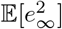according to

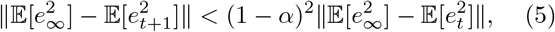

during any period of constant stochasticity and volatility (see SI S6). This exponential convergence means that traces of previous environmental conditions are quickly washed out. Consequently sublearner weights (which are calculated from these estimated errors) converge exponentially fast to reasonable values, as seen in the first few timesteps of Figure 2B. Remaining weighting inaccuracy is caused by fluctuations around the expectations in (5), as seen in later timesteps of Figure 2B.

The exponential convergence of Hetlearn weightings stands in contrast to the slow convergence of stochasticity/volatility estimates (Figure 2A and C), which result in poor initial performance of the Kalman learner. This slow convergence is due to the fact that stochasticity and volatility *explain each other away* [42]: a contribution from either noise term has the *same* effect on PE magnitude on any given trial. Only long-term patterns of PE autocorrelations can disambiguate the two, which is precisely how the ALS algorithm estimates them. More generally, good estimates of second-order statistics are inherently data-hungry to accurately estimate. Overall imprecise approximates of the stochasticity/volatility distributions (such as the point-estimate provided by ALS), are far worse in the low-data regime. Biological implementations of such Bayesian procedures, which are inevitably approximate, would likely suffer more.

### Competing sublearners should stay simple

We now strengthen the claims of Figure 1: that Hetlearn gains robustness to unmodelled environmental changes by assuming less of the environment. To do so, we compare Hetlearn against the algorithm proposed in [42], which we call the ‘Piray learner’. This belongs to a class of probabilistic algorithms known as particle filters, but was specifically formulated to address state inference under conditions of changing stochasticity and volatility. Like Hetlearn, they use parallel sublearners with different beliefs, weighted on predictive performance. Unlike Hetlearn, each sublearner estimates stochasticity and volatility: necessary for statistically optimal reward prediction. Specifically, each sublearner (/particle) is a Kalman Filter, with stochasticity/volatility estimates that stochastically diffuse to track changing environments.

We use a test environment similar to the previous section, in that the state follows Eqs. (2) and stochasticity/volatility are Gaussian ((4)). However, the variances of the (Gaussian) *C*_*m*_(*z*_*t*_) and *C*_*p*_(*z*_*t*_) change over time, following the schedules in Figure 3 (top row). These model a variety of large-world changes that can’t be predicted by the learner.

**Figure 3.**
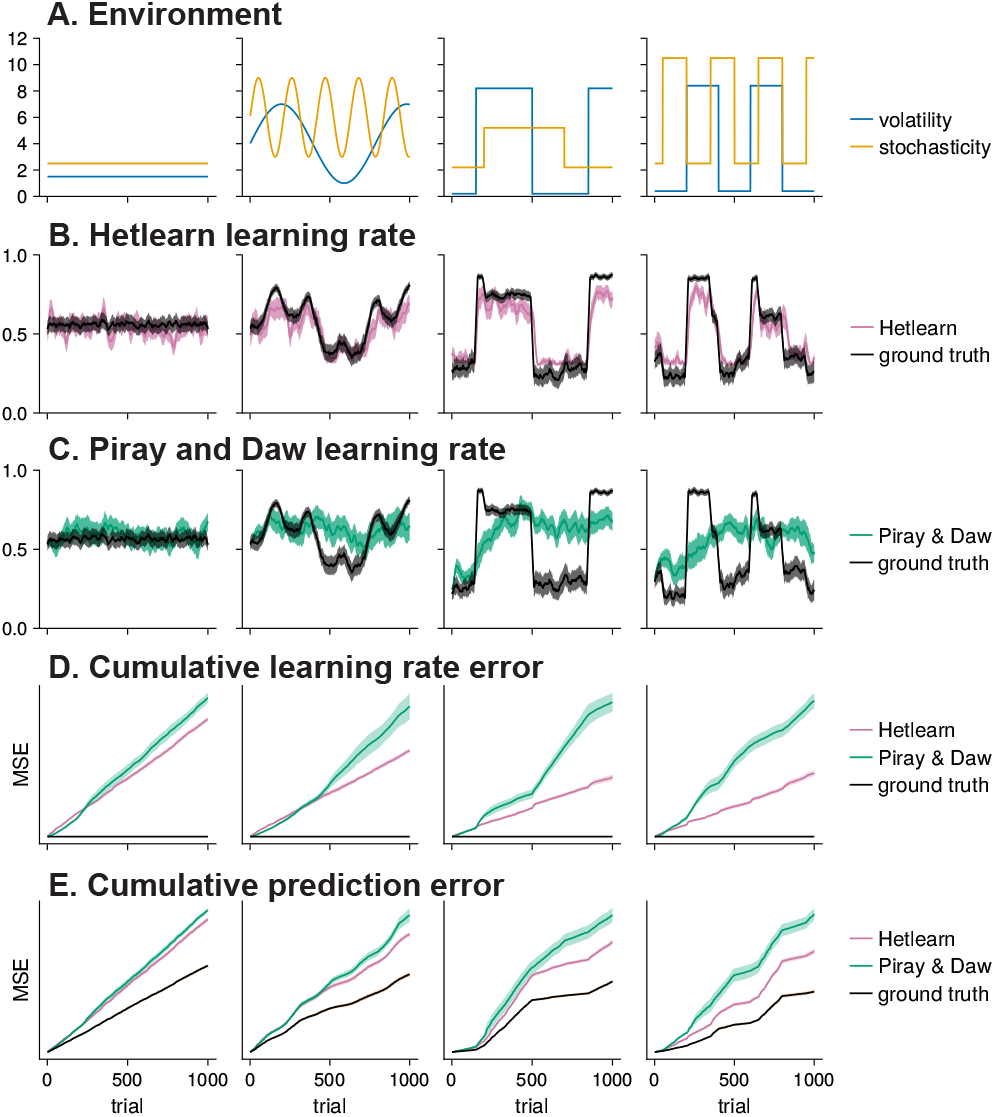
A: Volatility and stochasticity for four different environments not encompassed by a single generative model. B: Hetlearn (top) and Piray et al (bottom) tracking the the optimal learning rate across all four environments. C: Cumulative error over 10 trials for Hetlearn and Piray et al. D: Cumulative mean squared error (MSE) over 10 trials for Hetlearn and Piray et al. For B, C, and D, and the average of 10 runs were plotted for Hetlearn and Piray et al.

Hetlearn outperformed the Piray learner both in overall predictive performance (Figure 3E), and in tracking the correct learning rate over time (Figure 3D). Furthermore, Hetlearn used fewer sublearners (3, versus 10 for Piray).

The previously mentioned issues of stochasticity/volatility taking lots of data to estimate accurately *compound* in a situation where they can drift. Note that estimating the state as a probability distribution *requires* this estimation. Hetlearn, by avoiding it, cannot be reformulated as a probabilistic inference algorithm (see SI S4).

Inferring a hidden state subject to unmodelled changes that could be both first-order (i.e. from stochasticity and volatility) and second-order (i.e. from *changes* to stochasticity/volatility) is a generic problem. It is relevant for animals that incompletely model their environments due to environmental novelty or neural circuit limitations. Figure 3 demonstrates that chasing optimal probabilistic inference is not a sensible approach in these circumstances, and that Hetlearn is a simple, superior alternative.

### Relating Hetlearn to *Drosophila* learning circuitry

We now discuss the connection between the circuitry and physiology of the Drosophila Mushroom Body (MB) and Hetlearn (Figure 4C). This will enable us to test whether Hetlearn recapitulates known features of biological learning in the final section of the Results. The MB contains several compartments, which individually make predictions, and adapt their internal prediction errors with intrinsic, compartment-specific characteristics. Different compartments have different intrinsic learning rates, and are weighted dynamically. More detailed aspects of circuitry are considered in a subsequent section.

**Figure 4.**
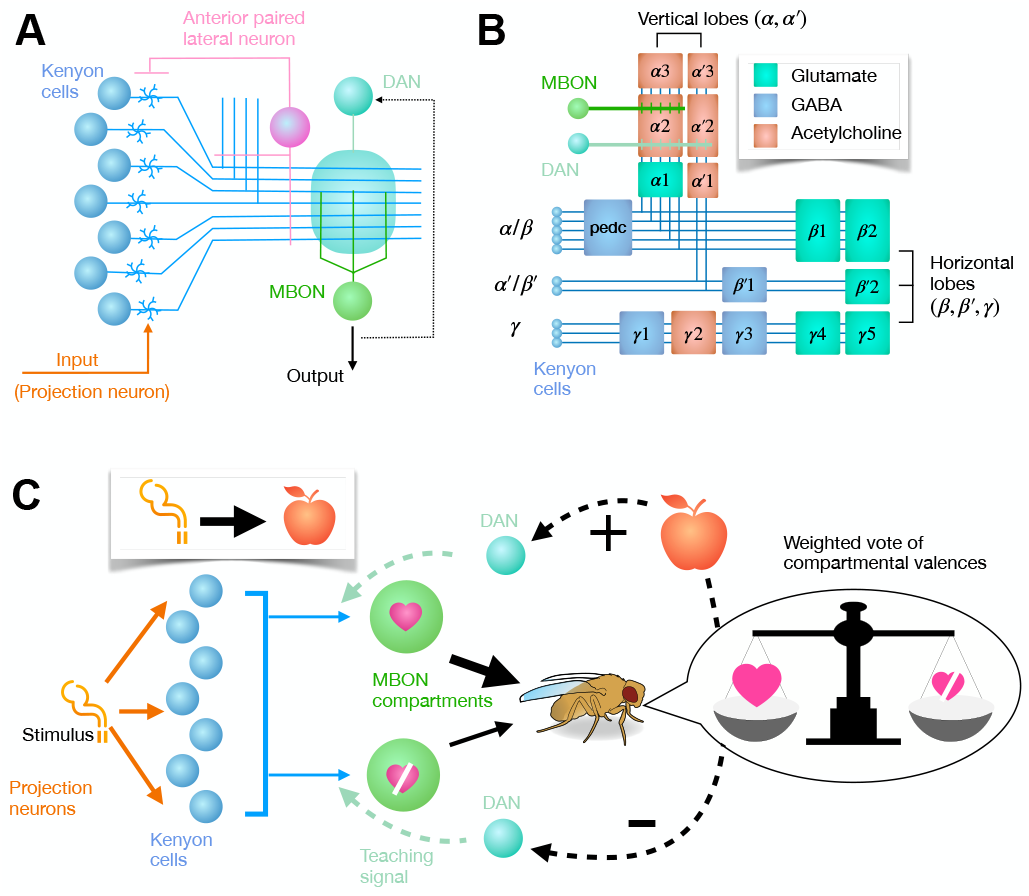
**A:** Schematic of an individual MBON compartment. The effect of sensory representations, transmitted through KC axons, on MBON activity, is adapted through plasticity induced by co-occurring DAN activation. Adapted from Figure 1A of [34]. **B:** Topology of the 16 compartments comprising the MB. KC axons arising from the three KC cell types (*α/β, α*^*′*^*/β, γ*) are depicted as blue lines, segregated according to anatomical layer. Example MBON and DAN included. Adapted from Figure 1C of [3] **C:** Schematic of proposed qualitative mechanism. Separate MBON compartments receive highly overlapping sensory representations from KCs. However, their valences are different, and are updated by anatomically and functionally distinct DAN teaching signals. Overall valence is a weighted sum of the individual compartments’ opinions.

Sensory representations in the MB are formed in the Kenyon Cell (KC) population which receives inputs from sensory projection neurons (Figure 4A). Tuning curves of the (≈ 2000) KCs are input-selective and sparse [50], suggesting an architecture optimised for decorrelating representations of similar sensory inputs [8, 28]. Conversely, KC outputs converge onto only 34 MB output neurons (MBONs), arranged in 16 compartments [3, 27] (Figure 4B).

MBON responses to similar sensory inputs, unlike KCs, are highly correlated [23]. Conversely, inputs that carry opposing valence (aversive vs appetitive) are well separated in MBON responses [23]. This suggests that MBON activity reflects the behavioral value of a stimulus rather than its low level features. Indeed, photoactivation/suppression of specific MBONs leads to attraction or avoidance behavior [40, 3].

Behavioral changes can occur via associative learning that updates the valence of a stimulus or set of behavioral states. These changes are driven by plasticity in the KC-MBON synapses, which is in turn induced by a ‘teaching signal’ from Dopaminergic Modulatory neuron (DAN) activity (Figure 4A,C) that communicates perceived reward or value.

The architecture of the MB *constrains* the abstract learning mechanisms its neural circuitry implements. Individual KC axons traverse and connect to multiple distinct compartments. In contrast, each of the 20 DAN types innervates only one or two compartments (Figure 4B). The instructive role of the DAN input to these compartments was demonstrated by [2], by photoactivating specific DANs, in conjunction with an olfactory stimulus, to induce appetitive conditioning. Subsequent behavioural changes could thus be attributed to a specific combination of teaching signals, compartments and stimuli, extracting the physiological chain of events leading to a conditioned stimulus (CS).

Differences in behaviour produced by activating different compartments were then compared, by observing the flies’ subsequent preference for the CS over time. The consequent mechanistic effects mirror the Hetlearn algorithm:

- Compartments can bi-directionally update the valences they attribute to conditioned stimuli, even through modulation of the same DAN (Unconditioned Stimulus / US).
- Compartments exhibited intrinsic heterogeneity in the degree and manner they updated their valence estimates of stimuli in response to new experiences. In particular, different compartments had intrinsically different learning rates, and held memories for different amounts of time.
- The contribution of the memories of different modules to overall behaviour is dynamic: they ‘vote’, but the importance of different voters changes over time. Co-activating DANs to the (appetitive) *α*1 and (aversive) *γ*1 *pedc* compartments with an odour resulted in an odour valence that was initially strongly aversive, and then appetitive after one day. This is consistent with the known feature of slower memory decay in the *α*1 compartment. Moreover, the initially strongly aversive response implied a nonlinear, dynamic summation of the compartmental valences.

### Hetlearn for complex conditioning experiments

So far we have considered how learning-rate parallelism aids inference of a single state from observations directly pertaining to the state. We now consider how adding parallelism on an alternative dimension aids learning when animals are faced with a continuous stream of stimuli. Here, a surprising outcome could be due to an as-yet unidentified pattern in preceding stimuli, or a fundamentally unidentifiable large-world effect. This difficulty is highlighted in three standard conditioning experiments (see Figure 6A,B,C), involving stimuli *S*_1_ and *S*_2_, a reward *R* and a punishment *P* :

**Extinction:** An initial association (e.g. {*S*_1_ → *R*} repeated), is subsequently extinguished ({*S*_1_ → ∅}repeated).

**Reversal (e.g. [57, 32]):** Flipping between repeated bouts of {*S*_1_ → *R*} and {*S*_1_ → *P* }.

**Second-order conditioning (e.g. [49]):** Initial Pavlovian conditioning (e.g. {*S*_1_→*R*}repeated) is followed by unrewarded presentation of a similar stimulus: ({*S*_2_ → *S*_1_ → ∅} repeated).

The abrupt environmental changes in extinction and reversal conditioning are due to unsensed large-world effects that hold for a limited (reversal) or indefinite (extinction) period. Conversely in second-order conditioning, prediction errors arise from an initially deficient predictive model: the animal initially has no way of knowing that *S*_2_ signals a context where *S*_1_ is unrewarded. Once this knowledge is acquired, a learner can perfectly predict rewards from stimuli.

Many experiments have shown that Drosophila cache adaptations to different large prediction errors in separate engrams, which each possess unique dynamics. We propose that this serves a purpose: the organism can ‘hedge their bets’ on the appropriate adaptation to prediction errors through parallelism. We now provide a Hetlearn model of MB that enacts this goal, while explaining organismal behaviour and making testable mechanistic predictions.

Our model first projects presented stimuli into a high-dimensional representation modelling the Kenyon-cell layer, denoted *x*_*t*_ at the *t*^*th*^ presentation. Representations decay on a slower timescale than stimulus presentation. Hence, the representation *x*_2_ after experiencing *S*_2_ → *S*_1_ holds traces of the *S*_2_ representation. Each representation *x*_*t*_ is normalised, e.g. by the recurrent inhibitory action of the APL neuron on the KC layer.

Our model learns to predict the current *x*_*t*_ by adapting a matrix *A* representing linear, recurrent Kenyon-cell connections. This adaptation follows the simple online-least-mean-squares rule, which can be implemented with Hebbian plasticity [53]:

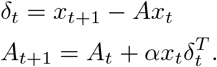

We use the learning rate *α* = 0.4. Biologically, structural changes in the calyx, presynaptic to Kenyon cells, do accompany the consolidation of olfactory conditioned memories [10, 4].

Conditioned valences are held in parallel engrams, which form in the KC-MBON connections. The *i*^*th*^ engram is defined by a ‘template’: *e*^(*i*)^ = *x*_*t*−1_, corresponding to the Kenyon cell representation that maximally activates it, and a binary effect *v*^(*i*)^, which is ±1 for an appetitive/aversive valence. The engram has fast and slow components held in different MBON compartments, and defined by intrinsic learning rates, which we set as *α*_*f*_ = 0.6, *α*_*s*_ = 0.2. Each component has a dynamic strength 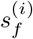 and 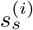, with overall strength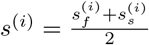.

On each stimulus presentation, the system makes a prediction *ŷ*_*t*+1_ of the next unconditioned stimuli. These are later compared against a vector of experienced unconditioned stimuli *y*_*t*_, represented in separate circuits such as the lateral horn. The prediction sums engram activations:

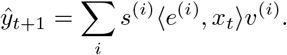

If the PE ∥*y*_*t*+1_ − *ŷ*_*t*+1_∥_2_ exceeds a threshold (*> T*_1_ = 0.3) and all extant engram activations ⟨*e*^(*i*)^, *x*_*t*_⟩ are low (*< T*_2_ = 0.3), then a new engram is formed with template *x*_*t*_. If the PE was an unsystematic fluke, this engram will not contribute constructively to future predictions and will die (described later). If not, it will crystallise without interfering with different existing engrams.

Engram strengths adapt to each PE. To minimise interference of each adaptation with different conditioned responses, engrams *compete* to claim and adapt to errors, a process we call *belief competition*. We take belief as

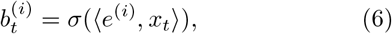

where 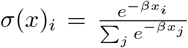is the softmax function. We took *β* = 10. Strengths then update as

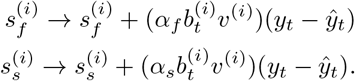

Slow strengths only update once the fast strength has crossed a threshold (*T*_*f*_ = 0.4) replicating the repeated conditioniong required to form engrams in slow-learning MBON compartments such as *α*1. If a fast strength dips below 0.1 before a slow strength has activated, the engram is extinguished.

In reinforcement learning, the value of an environmental state is the long-term predicted reward, which incorporates the reward obtained at future states likely to be visited. Correspondingly, our model values a KC representation *x*_*t*_ by evolving the representation through the recurrent matrix *A*, which predicts future representationas as 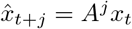. The synaptic output of these dynamics are integrated over time against different engrams, determining the engram-specific value contribution. Mathematically,

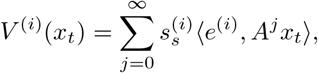

with overall value 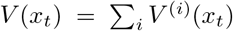. These dynamics soon die out biologically, and in the model (as ∥*A*^*j*+1^∥ *<* ∥*A*^*j*^ ∥), so we terminate the integration at *j* = 3.

Overall our model relies on simple computations: soft-max, correlation, and recurrent linear dynamics, which can all be enacted with biologically plausible learning rules.

### Hetlearn explains and recreates multiple behaviours and their mechanistic underpinnings

In Drosophila, extinction of a conditioned memory is supported by the acquisition of an entirely new memory towards the conditioned stimulus, which competes with the initial memory to govern extinction [17, 18]. Similarly, our model added a new, punishing engram at the onset of extinction (Figure 6C), whose increasing strength induced extinction. The original, rewarding engram is stably maintained meanwhile. As such, if extinction were a one-off fluke, or a temporary circumstance, it does not corrupt the original, rewarding memory.

Analogously, our model created four parallel memories during reversal conditioning protocols, as depicted in Figure 6B. The preference index of Figure 6 demonstrates how the speed of initial learning was governed by the slow/fast learning compartments within a context. Conversely, the speed of switching preferences during reversals was governed by competition *between* contexts, leading to changing degrees of conditioning across reversal blocks. The first reversal resulted in a weaker memory, while subsequent reversals built stronger memories, in agreement with experimental findings (e.g. Figure 1A of [56]).

Finally, our model recreated the behavioural dynamics observed in second-order conditioning.The valence of *S*_2_ initially rose due to an association formed with *S*_1_, whose representation formed the template of a rewarding context. A punishing context was formed to account for reward omission after *S*_2_ →*S*_1_, whose increasing strength accounted for slower-timescale extinction of this initially positive valence. These behavioural dynamics are highly conserved across animal phyla [21], and we provide a mechanistic and functional explanation.

We can test our model on mechanistic and behavioural data provided in [58], and contrast it against a different computational model of second-order conditioning provided in the same paper (Figure 5D) to capture parallel engram recruitment. There, a slow, rewarding compartment (MBON-*α*1) learns the first order association between *S*_1_ and reward. It then acts as a ‘teacher’ to ‘student’ compartments (*γ*5 and *β*^′^2*a*) that learn the second order association. The transience of this second-order memory therefore reflects the *intrinsic* fast learning/forgetting nature of these student compartments, rather than competition between engrams, as in our model, or in extinction condtioning. Teacher-to-student instruction occurs via the critical SMP108 interneuron, which forms part of a 2-hop connection recurrently connecting these two compartments. However, two key experimental observations are inconsistent with the *Yamada* student-teacher model.

**Figure 5.**
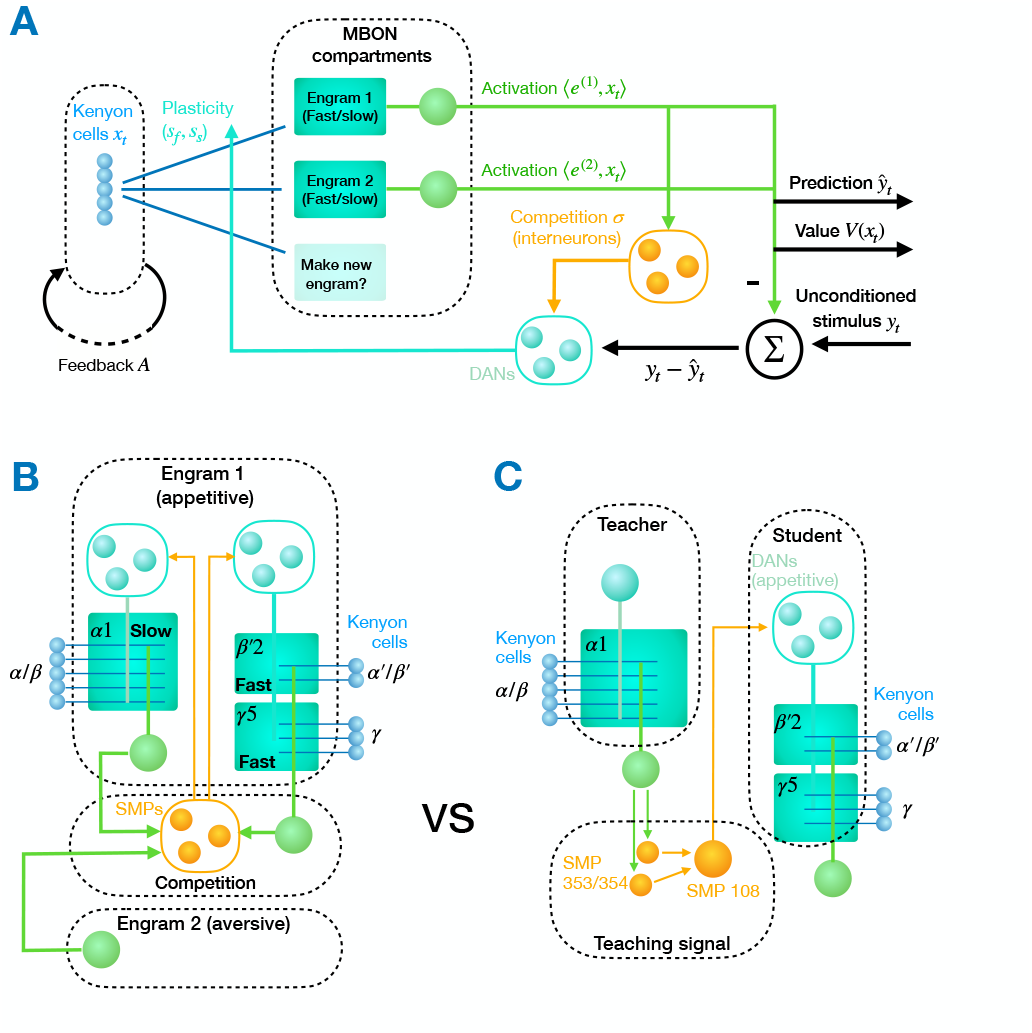
**A**: Hetlearn model for behavioural conditioning in Drosophila. Different engrams strengths compete to update from a prediction error, reducing interference with previously learned information. Predictions themselves do not invoke this competition, allowing the system to make rich, context-dependent predictions that make use of the representional capacity of multiple engrams. **B:** Proposed model of mechanism underlying second order conditioning. Compare to **C:** Mechanisms underlying second-order conditioning as proposed in [58]

First, the putative student compartments were able to act as teachers. The authors observed that coactivation of the DANs PAM-*γ*5 and PAM *β*^′^2*a* with the *S*_1_ stimulus implanted a first order memory (Figure 2B of [58]). Subsequent behavioural conditioning then allowed for a second order association. The flexibility of the underlying circuit is thus more consistent with bidirectional signalling between fast and slow learners, and also the connectivity of SMP108.

Secondly, the student-teacher model predicts *complete* elimination of second-order conditioning if the link between student and teacher is lost. However, optogenetic inactivation of the SMP108 interneuron halves, but doesn’t eliminate, second-order conditioning from the MBON-*α*1 implanted memory (Figure 6D, pane 3/4).

**Figure 6.**
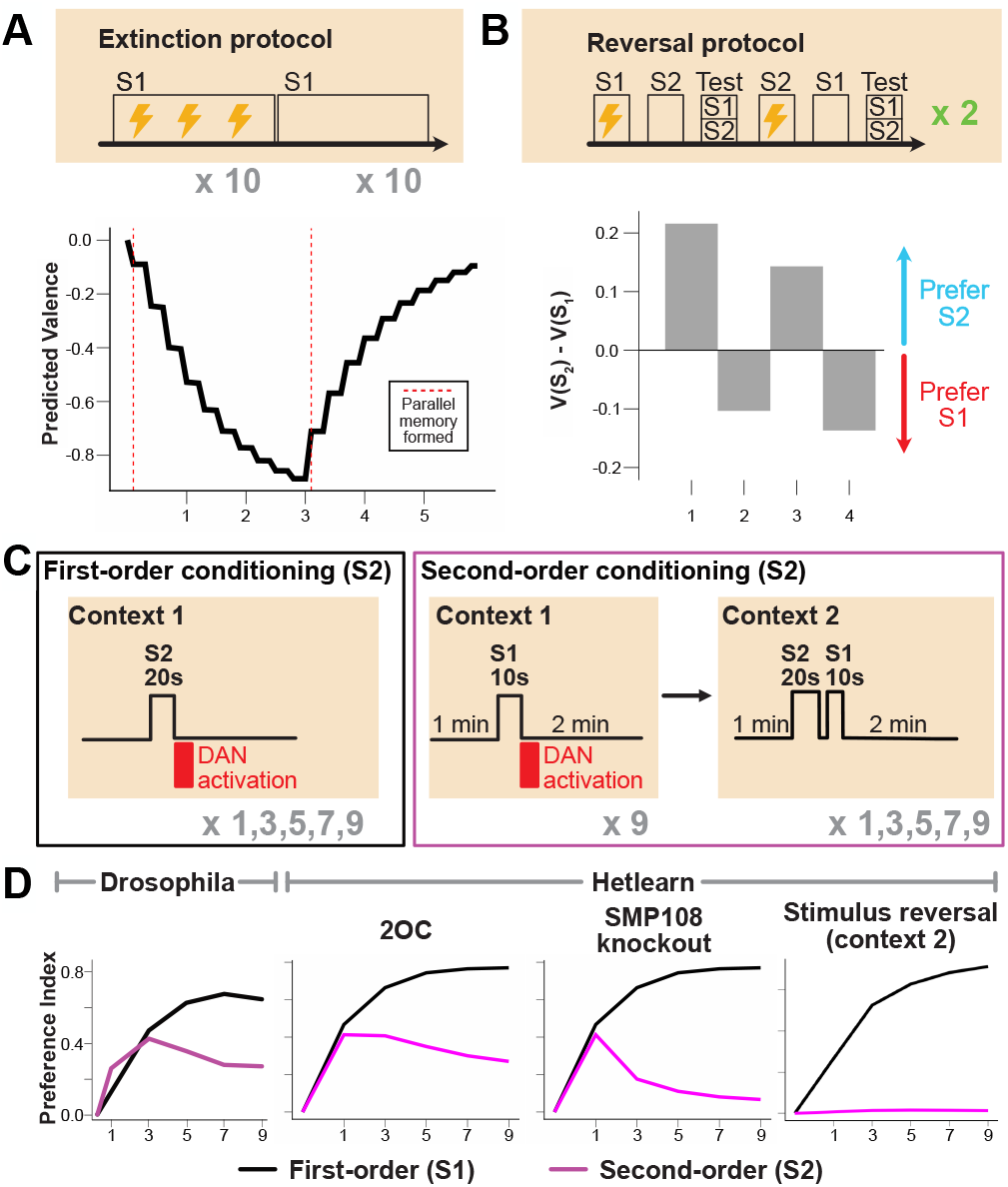
Model predictions vs behavioural experiments **A:** Extinction protocol adapted from [18]: *S*_1_ is associated with punishment (reward of *−*1) 30 times, followed by 30 punishment omissions. Stimulus preference of the model to *S*_1_ over time is graphed, along with the time at which the first (rewarding) and second (omitting) contexts are automatically formed in the model. **B:** Reversal protocol taken from [56]. Each block depicts ten pairings of *S*_1_ or *S*_2_ with punishment (reward of *−*1) or omission. Bar plot depicts difference between valences for *S*_1_ and *S*_2_ at the end of each block. **C**: Experimental protocol used in [58], and our model, for second-order conditioning. We modelled DAN activation as a reward on the timestep after the stimulus was presented. In context 2, *S*_2_ and *S*_1_ were presented on successive timesteps. **D**: *Drosophila* preference to *S*_1_ and *S*_2_ over time. Experimental data (Figure 1H of [58]) on leftmost pane, depicting preference index. Our model graphs valence on other panes, depicting (left) standard conditioning, (centre) conditioning under SMP108 knockout, and (right) conditioning when *S*_1_ is presented before *S*_2_ in context 2.

Our model requires belief competition across MBON compartments. We propose that SMP108 enacts this competition. This is consistent with its status as an anatomical hub for bidirectional connections between MBON compartments [58]. As such, we modelled SMP108 inactivation as the cessation of belief competition, by removing the softmax in Eq. (6). This halved second-order conditioning (Figure 6, Pane 3/4), consistent with experimental data.

Overall, while both models account for basic second-order conditioning, our alternative account of the role of SMP108 explains additional artefacts of behaviour and mechanism that are inconsistent with the *Yamada* model. It also makes distinct, testable predictions:

- SMP108 inactivation should have *less* of an effect on second-order conditioning during its early phases, when *S*_2_ →*S*_1_ → ∅ has been presented fewer times, and the strength of the punishing engram is thus weak.
- SMP108 inactivation should *reduce* the preference to *S*_1_ after a bout of second-order conditioning. The lack of belief competition increases the adaptation of the rewarding engram to reward omission after *S*_2_ → *S*_1_, thus reducing its strength more.

## Discussion

In this work we introduced a biologically motivated design principle, Hetlearn, for learning in scenarios where the causes of systematic environmental change can neither be represented, nor treated as noise. This provided a model of learning that recreates, interprets, and predicts Drosophila neurophysiology and behaviour higher-order conditioning experiments, without per-experiment tuning. Notably, Hetlearn trades asymptotic optimality for performance in low-data regimes and robustness to unmodelled, ‘Large world’ deviations in environmental statistics.

We first considered a fundamental component of learning across animal species: updating estimates of model states of the world through prediction errors. Existing work bases animal learning strategies on Kalman Filtering [25, 1], and Reinforcement learning [44, 37]. Here, the learning rate (i.e. sensitivity to prediction errors) is a key parameter determining estimation performance. Across tasks, both the optimal and actual values can vary dynamically. As such, neuroscientists have long considered how animals could approximate a Bayes-optimal learning rate [11, 25, 20].

In the simplest setting, two factors: stochasticity and volatility, must be estimated to set an optimal learning rate, and to represent the state as a probability distribution. Extensive literature proposes how this estimation might be performed across species [30, 33, 36, 41, 42, 47]. However, we showed that optimal learning strategies are fragile to two inevitable scenarios: drift in environmental stochasticity/volatility, and approximation error when estimating these factors from limited data.

We therefore propose that the costs of Hetlearn, both in terms of anatomical parallelism and asymptotic sub-optimality, are more than compensated by its robustness to unmodelled environmental change and its superior performance in the low-data regime.

State inference through parallelism is well known engineering principle, underpinning algorithms such as mixture-of-experts [24] and particle filtering [13]. Previous work proposed learning rate parallelism for the prediction error-based learning we considered [55]. A key difference in Hetlearn is that it avoids a specific generative model of environmental change (e.g. through abrupt, discontinuous changepoints, as in [55]) and instead incorporates recency bias in sub-learner weighting. This feature is shared with existing algorithms that deal with nonstationary large-world drift for classification problems [16] and means that weightings remain reactive to unmodelled environmen-tal changes indefinitely.

The basic Hetlearn algorithm justified the core idea of using parallelism to implicitly infer environmental quantities that are speedily constrained by data (i.e. learning rate to PEs). This is superficially consistent with detailed Drosophila MB circuitry, which displays intrinsically heterogeneous learning rates in compartments that learn behavioural valence from prediction errors. Future work could explore whether detailed circuit models strongly constrain circuitry to operate in a manner consistent with Hetlearn, though degeneracy and gaps in neurophysiological data would make this extremely challenging.

We proposed a simple, high-level model of MB learning that used parallelism to cache adaptations to surprising prediction errors before they could be confidently attributed to fluke or systematic, large-world environmental change. Our model recreated Drosophila behaviour (which is shared across phylla [21]) on key conditioning tasks where surprises arose either from fundamentally unpredictable large-world effects (extinction, reversal) or insufficient information on the relationship between observable stimuli (second-order conditioning), without requiring task-specific calibration. Given the emergence of these traits from a minimal model, we propose they reflect functional constraints on learning that enable animals to cope with truly novel situations.

Our model is qualitatively distinct from reinforcement learning algorithms, which are often used to describe conditioning [44, 48]. An RL-algorithm requires a preformed, fixed subdivision of the entire environment into behaviourally relevant states, whether these correspond directly to stimuli or emerge from pre-trained neural network [26, 51]. It then holds and updates values for each state. Our model instead spawns and removes states (i.e. engrams in the KC–MBON synapses) dynamically, after episodic experiences involving reward or punishment. This reflect parsimonious use of circuitry because there are fewer engrams to assign value to. It is also flexible, because it enables contextual modifiers of reward (such as *S*_2_ in second-order conditioning) on a biological timescale.

Many organisms, including mice and humans, share with Drosophila both behaviour on the conditioning tasks we considered, and the normative difficulty of flexibly building and acting on environmental representations in a truly novel environments. Our model ties these together through a simple, non-probabilistic algorithm exploiting parallelism for multiple purposes. While our algorithm had a plausible implementation in the mushroom body, the generality of the problems it mitigates and the behaviours it explains suggests its relevance to neural circuit computations across species.

## Methods

### Kalman Filter equations

The Kalman Filter updates valences from observed PEs *δ*_*t*_ according to the following equations:

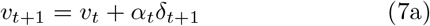

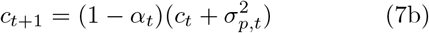

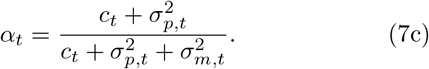

This requires seeding with 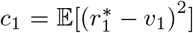: the initial covariance. Although ethologically inaccessible, we seeded all our simulations with the correct value of *c*_1_, giving an unfair advantage over Hetlearn.

This also requires seeding with 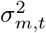 and 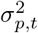 . In Fig-ure 2, they were assumed constant, and we used the ALS method for estimation. In Figure 3 they changed, and we used the Algorithm from [42] for inference. We describe these below:

### ALS Inference

ALS requires a separate learner that updates from PEs with a fixed learning rate *α* ∈ (0, 1). We set *α* = 0.5 in all simulations. So *v*_*t*+1_ = *v*_*t*_ + 0.5*δ*_*t*+1_.

A hyperparameter is the lag *j*, which we set at *j* = 10. A large lag requires a longer burn-in before inference can occur. A smaller lag reduces best-case inference accuracy.

Let *Y*_*t*_ = *r*_*t*_ − *v*_*t*_ be the reward estimation error of this ‘fixed learner’. Note the difference from the PE: *δ*_*t*+1_ = *r*_*t*+1_ − *v*_*t*_. Y_*t*_ is observable.

The ALS method builds estimates 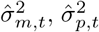 by solving the following regression problem:

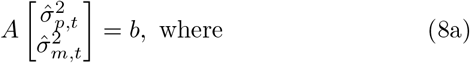

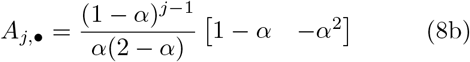

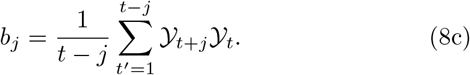

Here, *A*_*j*,•_ represents the *j*^*th*^ row of *A*, and *b*_*j*_ represents the sample estimate of E [*Y*_*t*+*j*_*Y*_*t*_].

For *t < j*, the ALS method cannot provide the Kalman learner with estimates. For these first 10 timepoints, we used *α* = 0.5. After, the Kalman learner updated according to the optimal equations of Eq. (7), using the ALS-derived estimates of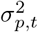 and 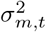 .

### Piray et al implementation

We implemented the particle filtering algorithm described in [42]. Each particle (/sublearner) updates valences according to the Kalman Filter (Eqs. (7)), but has private estimates of 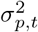and 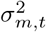 . We additionally provided each learner with the correct value of *c*_1_, as for the Kalman learner.

Private estimates of stochasticity and volatility update stochastically and identically. For volatility, the update equation is:

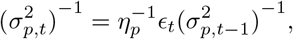

where

- *η*_*p*_ ∈ [0, 1] is a constant hyperparameter. Larger values indicate less change per trial.
- *ϵ*_*t*_ ∼*β*(0.5*η*_*p*_(1−*η*_*p*_)^−1^, 0.5) ∈ [0, 1] (a beta distribution)
- 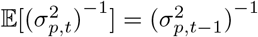.

The *i*^*th*^ particle calculates probabilities for the next reward by assuming

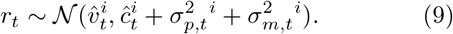

When an actual reward arrives, the relative weights of the particles are taken as

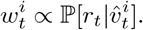

So particles whose probabilistic model predicted the actual reward with higher probability are weighted more. The weights are then normalised so that 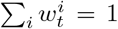. Thus the weights represent a probability distribution over particles: ℙ_*p*_, where the probability of the *i*^*th*^ particle is *w*_*i*_. The state of the overall particle collection is the expectation over this distribution:

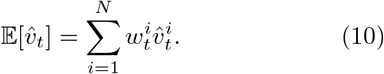

We follow [42] in using the Systematic Resampling procedure, to resample particles (deleting low-probability particles and duplicating high-probability particles) if their weight inequality exceeds 0.5.

## Code availability

All code for simulations in this work can be obtained at the public repositories: https://github.com/VRaman-Lab/HetlearnComplexConditioningModel.git (For Figures 1, 2, and 3) https://github.com/VRaman-Lab/HetlearnRewardPrediction.git

