## supplementary info for "Small circuit principles for learning in large worlds"

May 13, 2026

### S1 Truly optimal state inference

We consider a learner updating an internal state estimate $\hat{r}_t$ from observations $r_t$ on trial $t$. The state dynamics are:

$$\underbrace{r_t}_{\text{actual}} = \underbrace{r_t^*}_{\text{baseline}} + \underbrace{C_m(z_t)}_{\text{large-world fluctuation}} \tag{1a}$$

$$r_{t+1}^* - r_t^* = \underbrace{C_p(z_t)}_{\text{systematic large-world change}} , \tag{1b}$$

where $C_m$ and $C_p$ are unknown functions of countless large-world states $z_t$.

The estimation error is:

$$\underbrace{e_t}_{\text{estimation error}} = \underbrace{r_t^*}_{\text{baseline}} - \underbrace{\hat{r}_t}_{\text{estimate}} . \tag{1c}$$

A zero-error estimate would satisfy:

$$\begin{aligned}\hat{r}_{t+1} &= r_{t+1}^* \\ &= (r_{t+1}^* - r_t^*) + (r_t^* - \hat{r}_t) + \hat{r}_t \\ &= C_p(z_t) + e_t + \hat{r}_t.\end{aligned} \tag{2}$$

As such, the learner should ideally update its estimate by $C_p(z_t) + e_t$. However, these quantities are not directly observed, so this is impossible. Instead, the learner can only observe the prediction error (PE) $\delta_t$:

$$\begin{aligned}\delta_{t+1} &= r_{t+1} - \hat{r}_t \\ &= (r_{t+1} - r_{t+1}^*) + (r_{t+1}^*] - r_t^*) + (r_t^* - \hat{r}_t) \\ &= C_m(z_{t+1}) + C_p(z_t) + e_t.\end{aligned} \tag{3a}$$

We can express adaptation of the estimate to a PE as

$$\hat{r}_{t+1} = \hat{r}_t + \alpha_t \delta_{t+1}, \tag{3b}$$

where $\alpha_t$, the learning rate, quantifies the sensitivity of the estimate to a given PE. Equations (2) and (3) show that the ideal, unknowable learning rate is

$$\alpha_t = \frac{C_p(z_t) + e_t}{C_m(z_{t+1}) + C_p(z_t) + e_t}, \tag{4}$$

as this yields $\alpha_t \delta_{t+1} = C_p(z_t) + e_t$.

We emphasise that equation (4) is the **true** (but unknowable) optimal value of the learning rate on trial $t$. This differs from the Bayes-optimal learning rate, which depends on statistical assumptions on $C_m(z_t)$, $C_p(z_t)$, and $e_t$ that turns them into a quantifiable and inferrable small-world effect, and comprises the next section.

### S2 Bayes-optimal state inference (the Kalman Filter)

Suppose a learner makes the following assumptions on the problem of the previous section [4]:

- The prior distribution on the baseline reward, is

  $$r_1^* \sim \mathcal{N}(r_1, c_1),$$

  for some hyperparameter $c_1$, known *a priori*, which we call the initial covariance.

- The processes $C_m(z_t)$ and $C_p(z_t)$ have the following distributions:

  $$C_m(z_t) \sim \mathcal{N}(0, \sigma^2_{m,t}) \qquad\qquad C_p(z_t) \sim \mathcal{N}(0, \sigma^2_{p,t}),$$

  where $\sigma^2_{m,t}$ and $\sigma^2_{p,t}$ are hyperparameters known as stochasticity and volatility, respectively.

Then $r_t^*$ has the following probability distribution:

$$r_t^* \sim \mathcal{N}(\hat{r}_t, c_t),$$

where

$$\hat{r}_{t+1} = \hat{r}_t + \alpha_t^* \delta_{t+1} \tag{5a}$$

$$c_{t+1} = (1 - \alpha_t^*)(c_t + \sigma^2_{p,t}) \tag{5b}$$

$$\alpha_t^* = \frac{c_t + \sigma^2_{p,t}}{c_t + \sigma^2_{p,t} + \sigma^2_{m,t}}, \tag{5c}$$

and where $\delta_{t+1}$ is the PE defined in equation (3).

As such, a Bayes-optimal learner (known, in this linear environment, as a Kalman Filter) can update paired estimates of $\hat{r}_t$ and $c_t$ according to equations (5) to maintain an entire probability distribution for $r_t^*$. Effectively, the Kalman filter approximates the truly optimal learning rate of equation (4) with the approximant of equation (5c): the quantities $e_t$, $C_m(z_t)$, and $C_p(z_t)$ are replaced by their presumed variances.

The Kalman Filter has seen success in Engineering applications where the fixed-variance, white-noise assumption models e.g. sensor noise and thus is both reasonable, and amenable to being inferred offline using e.g. the Autocovariance Least Squares method [5]. Its fragility has long been acknowledged [CITES XXX SCHMIDT KALMAN FILTER]. Famously, the Kalman Filter with an optimal control policy (the 'linear-quadratic-gaussian') results in a closed-loop system that is infinitely sensitive. XXX.

Such fragility presumably carries over to Biology, where unknown environmental factors may not fit a Gaussian well, and certainly wouldn't have variances inferred ahead of time by the learner.

### S3 Summary of Hetlearn strategy for basic state inference

The Hetlearn strategy consists of parallel learning systems (sublearners) holding different predictions on latent, unobserved states. We discriminate between *inferred* latent states (such as $\hat{r}_t$, which sublearners update over time, and *guessed* latent states (such as learning rate $\alpha$), for which different sublearners have different, fixed, guesses.

Each sublearner makes a distinct prediction. Each sublearner is weighted based on an estimate of its recent predictive error. Recency is a hyperparameter, which we take as fixed, although it could also be a guessed latent state that is itself the subject of a Hetlearn strategy.

Let's start by denoting the state of a Hetlearner on trial $t$, and possessing $n$ sublearners. Elements with a subscript $t$ change over trials, while those with a superscript $(i)$ have distinct values possessed by the $n$ individual sublearners:

$$\mathcal{H}_t = \left\{ \tau, \{ \hat{e}_t^{(i)}, \alpha^{(i)}, \hat{r}_t^{(i)} \}_{i=1}^{n} \right\}.$$

Here, $\hat{e}_t^{(i)}$, $\alpha_t^{(i)}$, and $\hat{r}_t^{(i)}$ respectively denote the estimated error, learning rate, and state estimate of the $i^{th}$ sublearner on trial $t$, while $\tau$ is the recency bias parameter. Upon receiving an observed reward $r_{t+1}$, the $i^{th}$ sublearner updates as:

$$\hat{r}^{(i)}_{t+1} = \hat{r}^{(i)}_t + \alpha^{(i)}_t \delta^{(i)}_{t+1},$$
$$\hat{e}^{(i)}_{t+1} = \frac{1}{\tau}\|\delta^{(i)}_{t+1}\|^2_2 + \left(1 - \frac{1}{\tau}\right)\hat{e}^{(i)}_t.$$

where $\delta^{(i)}_{t+1} = r_{t+1} - \hat{r}^{(i)}_t$ is the $i^{th}$ PE. In other words, each sublearner has a fixed learning rate $\alpha^{(i)}_t$ which determines how its' state updates. Each sublearner also receives a PE, and its estimated error is a low-pass filtered summary of the PEs. The weights of each sublearner are inferrable from their estimated errors:

$$w^{(i)}_t \propto \frac{1}{e^{(i)}{}_t},$$

normalised so that $\sum_{i=1}^n w^{(i)}_t = 1$.

We could easily interpret a Hetlearn strategy as holding a posterior probability distribution on any such latent state. For instance, let's consider the state estimate $\hat{r}$ of the true baseline reward $r^*$. Each sublearner, on trial $t$, has weights $w^{(i)}_t$ and a state estimate $\hat{r}^{(i)}_t$. The *empirical* (/Dirac) distribution on $r^*$ is commonly used in particle filtering [2] as a probabilistic interpretation of a set of discrete predictions. This takes

$$P[r^*_t = z] = \sum_{i=1}^n w^{(i)}_t \mathbb{1}_{z - \hat{r}^{(i)}_t},$$

where $\mathbb{1}_x$ is the indicator function on $x$, which takes the values

$$\mathbb{1}_x = \begin{cases} 1 & x = 0 \\ 0 & x \neq 0. \end{cases}$$

This is a probability distribution, as the weights $w^{(i)}_t$ sum to one.

### S4 Hetlearn is qualitatively distinct from Bayesian Inference

In this section, we show how and why a Hetlearn agent does (and should) react to observed data in a manner distinct from an agent undertaking probabilistic inference. The non-mathematical summary is that

1. Bayes' theorem requires a generative model that separately models how the environment updates independently of observed data, and how observed data should tune this update (the predict and update steps of equations (9) and (8)). Hetlearn agents have neither. If they did, we would arrive at a logical contradiction.

2. Hetlearn assumes a partial environmental model where the predictive accuracy of a biased sublearner can change over time, based on unmodelled environmental factors. As such, sublearners are weighted based on *recent* (rather than overall) predictive performance. This is incompatible with probabilistic frameworks for inference and prediction based on sublearners. In these frameworks (known as sequential Monte Carlo methods/particle filters), the overall evidence for a given sublearner must be integrated over all time, without recency bias. This makes the Hetlearn mathematically incompatible with probabilistic inference.

We now illustrate these claims mathematically.

### S4.1 Single state inference on a Markov Process

We first consider the general problem where a hidden state $r_t^*$ (denoting baseline reward in the main paper) updates and gives rise to observed rewards probabilistically. We will derive the general Bayesian form for recursively predicting $r_t^*$ given previous rewards, and updating this prediction in response to new observed rewards $r_t$.

We will use $P(x)$ to denote the probability of an event $x$. We need the following assumptions:

$$r_{t+1}^*|r_t^* \sim f_t(r_{t+1}^*|r_t^*),$$

where $f_t(r'|r)$ is a probability density function denoting the probability density of changing from $r$ to $r'$ on trial $t$. Meanwhile the observed reward $r_t$, given the baseline, can be written

$$r_t|r_t^* \sim g_t(r_t|r_t^*),$$

where again $g_t(r'|r)$ is a probability density function denoting the probability of observing a reward $r'$ given a baseline reward $r$.

In fact for the setup of equations (1), $f_t(r'|r)$ is the probability density for $r' = r_t^* + C_p(z_t)$, given $r = r_t^*$, while $g_t(r'|r)$ is the probability density for $r' = r_t^* + C_m(z_t)$, evaluated at $r = r_t^*$.

This setup satisfies the *Markov property*, in that information on historical baseline rewards ($r_i^*$, for $i < t$, does not alter $f_t$ or $g_t$). Specifically:

$$P(r_{t+1}^*|r_t^*) = P(r_{t+1}^*|r_{1:t}^*)$$
$$P(r_t|r_t^*) = P(r_t|r_{1:t}^*).$$

Suppose that $r_1^*$ has some prior distribution encoded in a density function $\mu$, so: $P(r_1^* = x) = \mu(x)$. Then the probability of a sequence $r_{1:t}^*$ of unobserved baseline rewards, is

$$P(r^*_{1:t}) = \mu(r^*_1) \prod_{i=2}^{t} f_t(r^*_i | r^*_{i-1}). \tag{6a}$$

while the probability of a sequence of observed reward, conditioned on $r^*_{1:t}$, is

$$P(r_{1:t} | r^*_{1:t}) = \prod_{i=1}^{t} g_t(r_i | r^*_i). \tag{6b}$$

The laws of conditional probability then yield

$$P(r^*_{1:t} | r_{1:t}) = \frac{P(r_{1:t} | r^*_{1:t}) P(r^*_{1:t})}{P(r_{1:t})}, \tag{6c}$$

$$P(r^*_{1:t}, r_{1:t}) = P(r^*_{1:t}) P(r_{1:t} | r^*_{1:t}). \tag{6d}$$

where

$$P(r_{1:t}) = \int P(r^*_{1:t}, r_{1:t}) dr^*_{1:t}.$$

Equation (6a) represents the **prior** probability density of a sequence of hidden baseline rewards. Equation (6b) represents the **likelihood** of a sequence of observed rewards, given a sequence of baseline rewards. Finally, equation (6c) represents the **posterior** density of a hidden baseline sequence of rewards. These expressions can be modified into recursive forms that don't require storage of the entire past trajectories $r^*_{1:t-1}$ and $r_{1:t-1}$, We now demonstrate. First note that

$$P(r^*_{1:t}, r_{1:t}) = P(r^*_{1:t-1}, r_{1:t-1}) f_t(r^*_t | r^*_{t-1}) g_t(r_t | r^*_t).$$

Rearranging and using the laws of conditional probability, we get

$$P(r^*_{1:t} | r_{1:t}) = P(r^*_{1:t-1} | r_{1:t-1}) \frac{f_t(r^*_t | r^*_{t-1}) g_t(r_t | r^*_t)}{P(r_t | r_{1:t-1})}, \tag{7}$$

where

$$P(r_t | r_{1:t-1}) = \int \int f_t(r^*_t | r^*_{t-1}) g_t(r_t | r^*_t) \ \ dr^*_{t-1} \ dr^*_t.$$

Ethologically, an animal is unconcerned with the probability of past baseline rewards. As such, the quantity of interest is $P(r^*_t | r_{1:t})$. This can also be obtained recursively in a two-step process. First note that we can integrate out the trajectory $r^*_{1:t-1}$ in equation (7) to get

$$P(r^*_t | r_{1:t}) = \frac{P(r^*_t | r_{1:t-1}) g_t(r_t | r^*_t)}{P(r_t | r_{1:t-1})} \tag{8}$$

where

$$P(r_t^*|r_{1:t-1}) = \int f_t(r_t^*|r_{t-1}^*) P(r_{t-1}^*|r_{1:t-1})\ dr_{t-1}^*. \tag{9}$$

Equation (9) is known as the **prediction** step, which predicts the next baseline reward on trial $t$. Equation (8) is known as the **update** step, which updates the posterior probability of $r_t^*$ given the extra information of an observed reward $r_t$.

Note that the denominator $P(r_t|r_{1:t})$ is usually intractable to calculate, and purely dependent on data. As such, it is ignored, and the probabilities $P(r_t^*|r_{1:t-1})$ and $P(r_t^*|r_{1:t})$ are inferred only up to an unknown constant of proportionality.

A Bayes-optimal agent holds and updates a (scaled) probability distribution for the baseline reward using equations (8) and 9. Convenient forms for $f_t$, $g$, and $\mu$ are chosen so that updates are analytically tractable be s$v_t$ and $c_t$ update according to equations (5c).

Notice that this still requires knowledge of $\sigma_{m,t}^2$ and $\sigma_{p,t}^2$, the reward stochasticity and volatility respectively. To do probabilistic inference on these values is a separate process that would rely on a probability density for how they update, as explained in the main text.

#### S4.2 Hetlearn doesn't predict or update to a probabilistic model of observed data

An agent undertaking probabilistic inference must effect or approximate the predict and update steps of equations (9) and (8), for some choice of probability densities $f_t$ and $g_t$:

$$\underbrace{P(r_t^*|r_{1:t})}_{\text{update}} \propto P(r_t^*|r_{1:t-1}) g_t(r_t|r_t^*)$$
$$\underbrace{P(r_t^*|r_{1:t-1})}_{\text{predict}} \propto \int f_t(r_t^*|r_{t-1}^*) P(r_{t-1}^*|r_{1:t-1})\ dr_{t-1}^*.$$

For the setup of Equation (1) , recall that

$$g(r_t|r_t^*) = P[C_m(z_t) = r_t - r_t^*]$$
$$f(r_t^*|r_{t-1}^*) = P[p_t = r_t^* - r_{t-1}^*].$$

The Kalman Filter equations assumed that $P[C_m(z_t)] \sim \mathcal{N}(0, \sigma_{m,t}^2)$ and $P[p_t] \sim \mathcal{N}(0, \sigma_{p,t}^2)$.

Hetlearn doesn't specify a likelihood function $g$ or an update function $f$. In other words, it models neither $C_m(z_t)$ nor $C_p(z_t)$ as probability distributions. Indeed, the update of a given sublearner $i$, with learning rate $\alpha^{(i)}$, is Bayes-optimal with respect to a whole spectrum of update/likelihood functions $f$ and $g$. For instance, recall that under the Gaussian assumptions of the Kalman filter, the expected baseline reward updates Bayes-optimally as

$$v_{t+1} = v_t + \alpha_t \delta_{t+1}$$

where

$$\alpha_t = \frac{\sigma_{p,t}^2 + c_t}{\sigma_{m,t}^2 + c_t + \sigma_{p,t}^2}, c_{t+1} = (1 - \alpha_t)(c_t + \sigma_{p,t}^2).$$

As such, a given learning rate $\alpha^{(i)}$ is Bayes-optimal with respect to multiple values of $\sigma_{m,t}^2$, $\sigma_{p,t}^2$, and $c_1$. However, different values of $\sigma_{m,t}^2$, $\sigma_{p,t}^2$ and $c_t$ imply different likelihood functions, state update functions (i.e. $f$) and prior reward distributions, respectively.

Overall, a sublearner with given fixed learning rate $\alpha^{(i)}$ is incompatible with a single posterior probability density for $r^*$, on the prediction (or update) step. As such, it cannot assign a specific probability to any choice of $r^*$.

Similarly, a weighted ensemble of sublearners, with different fixed learning rates and weightings, cannot imply a single posterior probability density for $r^*$, since each individual sublearner's probability for $r^*$ is ill-defined.

### S4.3 Difference between Sequential Monte Carlo methods (particle filters) and Hetlearn

We start by describing the class of sequential Monte Carlo methods for inferring $P(r_t^*|r_{1:t})$. The goal of sequential Monte Carlo (SMC) methods is to approximate and update (over trials) target probability densities $\pi_t(r_{1:t}^*)$ using multiple sublearners with different beliefs on $\pi_t$.

Suppose that $\pi_t(r_{1:t}^*)$ is the probability density for $P(r_{1:t}^*|r_{1:t})$. Then

$$\pi_t(r_{1:t}^*) = \frac{\gamma_t(r_{1:t}^*)}{Z_t},$$

where

$$\gamma_t(r_{1:t}^*) = P(r_{1:t}^*, r_{1:t})$$
$$Z_t = \int \gamma_t(r_{1:t}^*)\ dr_{1:t}^*.$$

Importantly, $\gamma_t$ (unlike $\pi_t$) can be evaluated pointwise, given known likelihood and update functions:

$$\gamma_t(r^*_{1:t}) = P(r^*_1)g[r_1|r^*_1]\prod_{i=1}^{t-1} f[r^*_{i+1}|r^*_i]g[r_{i+1}|r^*_{i+1}].$$

Suppose we could take $N$ samples of the baseline reward trajectory from the true probability distribution $\pi_t$. The $i^{th}$ sample will be denoted $R^i_{1:t}$. Then the empirical density $\hat{\pi}_t$ forms an unbiased approximation of the true density $\pi_t$ :

$$\hat{\pi}_t(r^*_{1:t}) = \frac{1}{N}\sum_{i=1}^{N} \mathbb{1}_{r_{1:t}}(R^i_{1:t}).$$

Essentially, this assigns a probability of $\frac{1}{N}$ to any baseline reward trajectory that exactly matches one of the $N$ samples $R^i_{1:t}$.

In particular this gives an unbiased estimate of the expected value of any function $\psi$ of the baseline reward [3]:

$$\begin{aligned}\mathbb{E}[\psi(r^*_{1:t})] &= \int \psi(r^*_{1:t})\pi_t(r^*_{1:t})\ dr^*_{1:t} \\ &\approx \frac{1}{N}\sum_{i=1}^{N} \psi(R^i_{1:t}).\end{aligned}$$

The empirical distribution makes it easy to approximate a marginal distribution such as $P[r^*_t|r_{1:t}]$, as:

$$\begin{aligned}P[r^*_t|r_{1:t}] &\approx \int \hat{\pi}(r^*_{1:t})\ dr^*_{1:t-1} \\ &= \frac{1}{N}\sum_{i=1}^{N} \mathbb{1}_{r^*_t}(R^i_t).\end{aligned}$$

As such, the expected baseline reward is unbiasedly estimated as

$$v_t = \mathbb{E}[r^*_t|r_{1:t}] = \frac{1}{N}\sum_{i=1}^{N} R^i_t.$$

Unfortunately, in the setup of equation (1) it is intractable to make samples (i.e. sublearners/particles) of the true target density $\pi(r^*_{1:t})$. Instead one uses **importance sampling**, which substitutes the true density with a proposal density $q$ that can be easily sampled. Formally, suppose we have a probability density $q$ that satisifes

$$\pi_t(r^*_{1:t}) > 0 \Rightarrow q(r^*_{1:t}) > 0.$$

Note that

$$\pi_t(r^*_{1:t}) = \frac{w_t(r^*_{1:t})q(r^*_{1:t})}{\tilde{Z}_t},$$

where

$$w_t(r^*_{1:t}) = \frac{\gamma_t(r^*_{1:t})}{q_t(r^*_{1:t})} \tag{10}$$

$$\tilde{Z}_t = \int w_t(r^*_{1:t})q_t(r^*_{1:t})\ dr^*_{1:t}.$$

Just like $\gamma_t$ previously, $w_t$ can be explicitly evaluated for a given trajectory $r^*_{1:t}$.

If we now generate sampled baseline reward trajectories: $\{R^i_{1:t}\}^N_{i=1}$ from $q$, i.e. $R^i_{1:t} \sim q(r^*_{1:t})$, then we can make a new empirical density that approximates $\pi$:

$$\hat{\pi}_t(r^*_{1:t}) = \frac{1}{N}\sum_{i=1}^{N} W^i_t \mathbb{1}_{R^i_{1:t}}(r^*_{1:t})$$

where

$$W^i_t = \frac{w_t(R^i_{1:t})}{\sum_{i=1}^N w_t(R^i_{1:t})}. \tag{11}$$

$W^i_t$ represent the weights of the $i^{th}$ sublearner, while $R^i_{1:t}$ represents its sequence of predictions. The state estimate

$$\mathbb{E}[r^*_t|r_{1:t}] = \frac{1}{N}\sum_{i=1}^{N} W^i_t R^i_t \tag{12}$$

is unbiased in the asymptotic limit as $N \to \infty$ (see [3]). In sequential importance sampling, sublearners are updated and reweighted recursively. Formally, suppose $q$ satisfies

$$q(r^*_{1:t}) = q(r^*_1)\prod_{i=2}^{t} q(r^*_i|r_{1:i-1}).$$

Then we can rewrite

$$w_t(r^*_{1:t}) = \frac{\gamma_t(r^*_{1:t})}{q_t(r^*_{1:t})} \tag{13a}$$

$$= \frac{\gamma_t(r^*_{1:t-1})}{q_t(r^*_{1:t-1})} \frac{\gamma_t(r^*_{1:t}) q_{t-1}(r^*_{1:t-1})}{q_t(r^*_{1:t})\gamma_{t-1}(r^*_{1:t-1})} \tag{13b}$$

$$= w_{t-1}(r^*_{1:t-1}) L_t(r^*_{1:t}), \tag{13c}$$

where

$$L_t(r^*_{1:t}) = \frac{\gamma_t(r^*_{1:t}) q_t(r^*_{1:t-1})}{q_t(r^*_{1:t})\gamma_{t-1}(r^*_{1:t-1})}$$

$$= \frac{\gamma_t(r^*_t | r_{1:t-1})}{q_t(r^*_t | r^*_{1:t-1})} \tag{13d}$$

Overall, a (sequential) importance sampling method holds $N$ sublearners representing samples from a proposal probability distribution from which marginals and functions of the target distribution $\pi(r^*_{1:t}) = P[r^*_{1:t} | r_{1:t}]$ can be approximated (e.g. equation (12)). Key points of note are

- The weights $W^i_t$ of the $i^{th}$ sublearner are proportional to the total probability of the sublearner's reward history over time (equations (10) and (11)).
- The weights are recursively updated by a multiplicative factor that is **independent** of past rewards (equation (13d)).

These key points are irrespective of the proposal density $q$, and the probability density for observed and baseline rewards: $\gamma_t(r^*_{1:t}) = P[r^*_{1:t}, r_{1:t}]$. Is Hetlearn a sequential Monte Carlo method, for some implicit choice of these densities? No: it does not satisfy these key points, as we now show.

First recall the Hetlearn update for the weights of sublearners:

$$\hat{e}^{(i)}_{t+1} = \frac{1}{\tau} \|\delta^{(i)}_{t+1}\|^2_2 + \left(1 - \frac{1}{\tau}\right) \hat{e}^{(i)}_t$$

$$W^i_t \propto \frac{1}{e^{(i)}_t}.$$

Rearranging the above equations, we get

$$W^i_t \propto W^i_{t-1} \times \frac{1}{(1-\gamma) + \gamma \delta^{(i)}_t W^i_{t-1}}, \tag{14}$$

where $\gamma = \frac{1}{\tau}$.

If Hetlearn were a sequential Monte Carlo method, the weights of the $i^{th}$ particle would satisfy equation (13). So

$$W_t^i = W_{t-1}^i L_t(R_{1:t-1}^i).$$

As such,

$$\begin{aligned}\frac{1}{(1-\gamma) + \gamma\delta_t^{(i)} W_{t-1}^i} &= L_t(R_{1:t-1}^i) \\ &= \frac{\gamma_t(R_t^i | R_{1:t-1}^i)}{q_t(R_t^i | R_{1:t-1}^i)}.\end{aligned}$$

However, the RHS of this equation is conditional on $R_{1:t-1}^i$ whereas the LHS is dependent on $R_{1:t-1}^i$. Hence they are incompatible for any choice of $\gamma_t$ and $q_t$.

In plain English, the weight update in Hetlearn incorporates a recency bias, and the weights therefore reflect recent sublearner performance. As such, the weight of the $i^{th}$ sublearner can't be proportional to the probability of the sublearner's predictions being true over **all** time.

### S5 Robustness of Hetlearn to changing recency bias

In this section, we consider the recency bias hyperparameter used in estimating the learning rate for Hetlearn in the main paper. We discuss its necessity in nonstationary learning problems, and demonstrate (through simulation) how accurate learning rate tracking is robust to a wide range of recency biases.

Hetlearn, unlike probabilistic inference, forgoes a complete generative model of the world incorporating the sources of change in the environment. As such, unmodelled processes could affect the environment, rendering old information less salient than new in inferring the current state of the world. This is why Hetlearn requires an extra recency bias hyperparameter, specifying the rate at which new information replaces old. Experimental evidence suggests that recency bias is set biophysically in the compartments of the Drosophila Mushroom Body, and is heterogeneous across compartments. This was demonstrated in e.g. [1] by monitoring the decay rate of memories implanted through Pavlovian conditioning experiments.

A shorter recency bias allows a learner's weightings to more quickly recalibrate to environmental change. In the absence of suc change, a longer recency bias is better as it exploits more data to make inferences on the state of the world.

In the main paper, we proposed that the competition between Hetlearn sublearners was robust to different recency biases. This is because learning rate is a parameter that is tightly constrained by each new PE: it is 'data-lean'. As such, forgoing historical data doesn't greatly compromise best-case weighting peformance.

Figure S1 demonstrates the performance of Hetlearn, simulated as in Figure 5 of the main paper, for different choice of recency bias and different nonstationary environments. This absolute performance is measured as the cumulative mean squared error over 1000 trials. The relative performance, plotted, is measured as the absolute performance relative to the absolute performance of the Piray & Daw algorithm described in Figure 4 of the main paper. Note that simulations with high learning rate error for Hetlearn do not necessarily lead to high MSE in state estimation. This is because the effect of over-estimating learning rate on PEs is much smaller than the effect of under-estimating, and Hetlearn tends to the former.

### S6 Exponential convergence of Hetlearn sublearners estimation errors

The robustness to changing recency bias is caused by the fact that *little* data is required to get a reasonably accurate weighting. Why? The state estimation error for a single sublearner, with fixed learning rate $\alpha$, on the task introduced in (1) updates as

$$\begin{aligned} e_{t+1} &= v_{t+1} - r^*_{t+1} && (15)\\ &= v_t + \alpha(r^*_t + C_p(z_t) + C_m(z_t) - v_t) - (r^*_t + C_p(z_t)) && (16)\\ &= e_t(1-\alpha) + K_t, && (17) \end{aligned}$$

where $K_t = C_p(z_t)(1-\alpha) + C_m(z_t)\alpha$. Assuming uncorrelated cross-terms, this means that

$$\mathbb{E}[e^2_{t+1}] = \mathbb{E}[e^2_t](1-\alpha)^2 + \mathbb{E}[K^2_t]. \qquad (18)$$

This recursive update satisfies the Banach fixed-point theorem, with Lipschitz constant $q = (1-\alpha)^2$. As such, consider a limited period of time where $\mathbb{E}[K^2_t]$ stays constant. For this constant value, there will be a unique fixed point $\mathbb{E}[e^2_\infty]$, with

$$||\mathbb{E}[e^2_\infty] - \mathbb{E}[e^2_{t+1}]|| < q||\mathbb{E}[e^2_\infty] - \mathbb{E}[e^2_t]||.$$

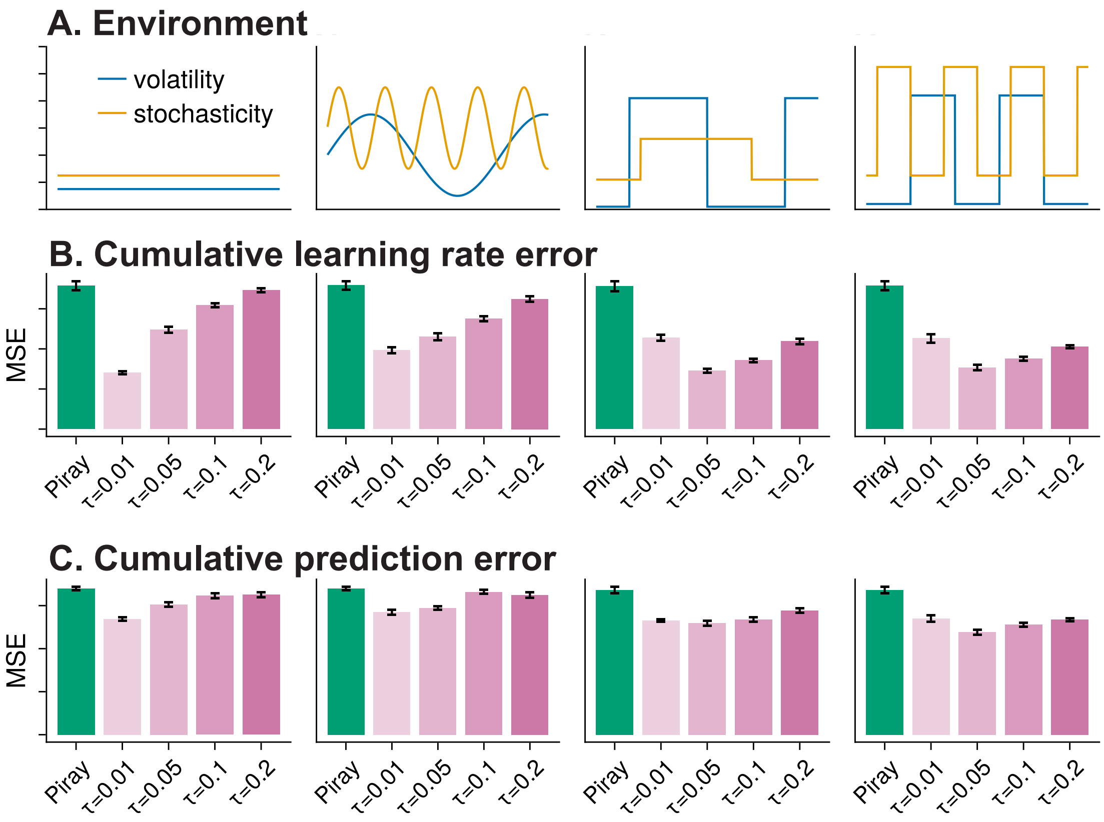


Figure S1: A: Time course of volatility of stochasticity over four different environments with different generative models. B: The cumulative learning rate error between the true and optimal learning rates after 1000 trials for Piray et al and Hetlearn with five values of the recency bias ($\tau$) spanning an order of magnitude. Ten repeats were taken for each datapoint. C: The mean squared error (MSE) for Piray et al and Hetlearn. Ten repeats were taken for each datapoint.

So the expected MSE converges exponentially fast to its steady-state value, with exponent dependent on learning rate. If a large-world change modifies the statistics of $K_t$, the expected MSE will converge exponentially fast to the new steady-state value. Of course, there is still a discrepancy between the actual MSE and its expected value. Hetlearn integrates successive prediction errors over time with a recency bias to suppress these fluctuations, analogously to a moving average.
